# Phylogeography of the electric rays of the family Narcinidae (Gill, 1862): systematic review and evidence of the dispersal routes to America

**DOI:** 10.1101/2024.11.04.621869

**Authors:** Luis Fernando da Silva Rodrigues-Filho, Richard Klein Castro Silva, Eduardo Lopes de Lima, Raquel Sicca-Ramirez, Getulio Rincon, Jorge Luiz Silva Nunes, Iracilda Sampaio, João Braullio de Luna Sales

**Author notes:** **Corresponding author:** Luis Fernando da Silva Rodrigues Filho *Federal Rural University of Amazônia, Group for Marine and Coastal Studies in the Amazon (GEMCA)*, *Capanema Campus, 68600-030 Capanema, Pará, Brazil*. E*-mail:*.

## Abstract

The electric rays belonging family Narcinidae are a cosmopolitan group, which has come under increasing scrutiny in relation to the taxonomic composition of the family and its phylogenetic position within the order Torpediniformes. Phylogeographic inferences can provide important insights into the present-day distribution of the narcinid species, as well as the systematics of the group. In this context, the present study evaluates the evolutionary and biogeographic history of the family Narcinidae, inferring distribution patterns and the molecular systematics of these organisms based on the DNA barcode. The results of the analysis indicate that the electric rays originated in the Central Indo-Pacific realm, with all other dispersal events beginning from this region. Two different dispersal routes were observed for the American narcinids. The first route was from the Central Indo-Pacific realm to the Pacific Ocean and then the Caribbean Sea, prior to the closure of the Isthmus of Panama. The second route start on Central Indo-Pacific realm passed South Africa continent before crossing the Atlantic. The principal torpediniform lineages diversified between the end of the Cretaceous and the early Paleocene, with the present-day narcinid taxa arising and diversifying between 25 and 20 Mya. The present study confirms the lack of monophyly in the Narcinidae, validates the genus *Narcinops*, and restricts the occurrence of the genus *Narcine* to the coastal environments of the Americas, which emphasizes the need for the review and revalidation of the synonym *Syrraxis* for the narcinids of the Central/Western Indo-Pacific realm. This study also describes additional lineages for *Narcine entemedor* and *Narcine maculata* and validates the occurrence of three species (*Narcinesp*., *Narcine brasiliensis*, and *Narcine bancroftii*) in the Atlantic Ocean, based on the published data, and emphasizes the need for a definitive description of *Narcine* sp.

## INTRODUCTION

Environmental factors, reproductive isolation, and sexual selection are all important drivers of speciation events, which can determine the appearance of a new species over time (Orr and Smith 1998). The interpretation of speciation events and the available species delimitation methods are often subject to considerable controversy, whether from the perspective of population biology or systematics, given the diversity of variables that drive speciation processes, and the relatively diffuse limits established, in many cases, which result in contradictory interpretations, depending on the method used (Carstens et al. 2013; Luo et al. 2018). The efficiency of the delimitation of species may be influenced by a range of factors, including the geographic distance between the sampling points, the number of haplotypes recovered, the difficulty of identifying species reliably based on their morphology, and the taxonomic classification of the data analyzed (Carstens et al. 2013; Luo et al. 2018). From an evolutionary perspective, phylogeographic studies can provide valuable insights for the understanding of the demographic processes that mold the evolution of populations and the emergence of new species due processes that regulate the geographic distribution of genetic lineages (Sites and Marshal 2003; Yang and Rannala 2012; Sales et al. 2019; Magoga et al.2021; Gales et al. 2024; Sales et al. 2024).

The available evidence indicates that the rays of the superorder Batoidea first appeared in coastal environments, and then dispersed to deeper waters, where they diversified (Aschliman et al. 2012). The batoid species of the family Narcinidae Gill, 1862 have a cosmopolitan distribution (Carvalho et al. 1998; Last et al. 2016). This family is composed by benthic rays, which are found in shallow coastal environments, where vicariant events and dispersal processes have resulted to the diversification of the ancestral forms into the modern narcinid species. One of the major events of Elasmobranchii diversification occurs during the Cretaceous period know as Cretaceous-Paleocene transition (K/Pg), a period marked by rapid radiations in the taxa that survived the mass extinction event (Aschliman et al. 2012; Müller et al. 2016; Villalobos-Segura and Underwood 2020).

Like the other families of the order, the phylogenetic position of the Narcinidae in the Toperdiniformes has been amply discussed in the scientific literature (Aschliman et al. 2012; Naylor et al. 2012; Claeson et al. 2014; Last et al. 2016; Melis et al. 2023), with many studies emphasizing the phylogenetic proximity of the Narcinidae and the Narkidae Fowler, 1934, which supports the need for more detailed analyses, given the lack of monophyly observed in these two families (Naylor et al. 2012, Last et al. 2016). In fact, the relationships among the narcinid genera are still poorly resolved. Melis et al. (2023), recovered the genera *Discopyge* Heckel, 1846, *Benthobatis* Alcock, 1898, and *Typhlonarke* Waite, 1909 in the same clade, which highlights the phylogenetic proximity of the Narcinidae and Narkidae. However, while Last *et al*. (2016) also recovered the *Benthobatis*/*Typhlonarke*/*Discopyge* clade in their phylogeny, it was in a sister-clade arrangement with Toperdinidae Henle, 1834, and *Narcinops* Whitley,1940.

The Narcinidae includes 29 valid species distributed in five genera (*Discopyge*, *Benthobatis*, *Diplobatis*, *Narcinops*, and *Narcine*). *Narcine* is the genus with the largest number of species – 15 – which are distributed in the western Atlantic, Central/Western Indo-Pacific, and eastern Pacific oceans. The only three *Narcine* species found in the western Atlantic were previously included in a single species, *Narcine brasiliensis* (Olfers 1831), although the systematic review of Carvalho (1999) indicated that this species is, in fact, composed of three distinct lineages, *N. brasiliensis*, *Narcine bancroftii* (Griffith and Smith 1834), and *Narcine sp.*, differentiated by their geographic and morphological characteristics. Each of these three species has a well-defined distribution in the western Atlantic. While *N. bancroftii* and *N. brasiliensis* are the only valid species in this group, they are morphologically very similar, being differentiated only by the additional rows of teeth found in *N. brasiliensis*, and slight differences in the coloration patterns (Carvalho 1999; Last et al. 2016).

With the advent of molecular techniques, a lot of questions on the phylogenetic arrangement of the elasmobranchs, their areas of occurrence, and the presence of cryptic lineages have been investigated over the past 20 years, and many studies have supported the revalidation of the elasmobranch families (Richards et al. 2009; Last et al. 2016; Castilho-Paez et al. 2017; Rodrigues-Filho et al. 2023). Even so, some elasmobranch groups have yet to have been assessed in detail, which may be obscuring the existence of some internal speciation events and, in turn, underestimating the true biological diversity of the group (Gales et al. 2024). Up to now, few studies have applied molecular techniques to the analysis of the members of the family Narcinidae, and even those that have, have focused on a small number of genera/species, with a limited sample (Gaitan-Espitia et al 2016; Kumar and Parakash 2023; Sureandiran et al. 2023), especially for the American lineages, such as the southern West Atlantic.

One of the principal factors associated with speciation in aquatic organisms are historical biogeographic events, which not only influence contemporary diversity, but also the current distribution of the species (Underwood 2006; Aschliman et al. 2012; Müller et al. 2016; Villalobos-Segura and Underwood 2020). The prevailing historical consensus on the emergence and diversification of the modern narcinid lineages focuses on the Asian western Indo-Pacific region (McEachran and Aschliman 2004; Daly-Engel et al. 2012; Sales et al. 2019), although some evidence indicates that this region was not the only center of origin and dispersal of some elasmobranch groups (Gales et al. 2024). Given this, more systematic phylogeographic inferences on the family should contribute to the understanding of the probable processes of speciation that occurred within the group and facilitate the interpretation of the current distribution of the narcinid species, and how biogeographic events happened to result current lineages. In this context, the present study investigated the evolutionary and biogeographic history of the family Narcinidae and inferred the factors that have driven the distribution patterns and molecular systematics of the family, as well as assessing the diversification and divergence of these organisms.

## MATERIALS AND METHODS

### Collection of Samples and Molecular Methods

For the present study, 20 samples of *Narcine* species were collected from different points along the Atlantic and Pacific coasts of South America – Amapá and Pará in Brazil (representing the Atlantic coast) and Tumbes in Peru (East Pacific). The specimens were captured as bycatch and by artisanal fisheries (Table S1; Fig. 1). Each specimen was identified initially following the key of Last et al. (2016), and a sample of muscle tissue was collected and stored in absolute ethanol.

**Fig. 1.**
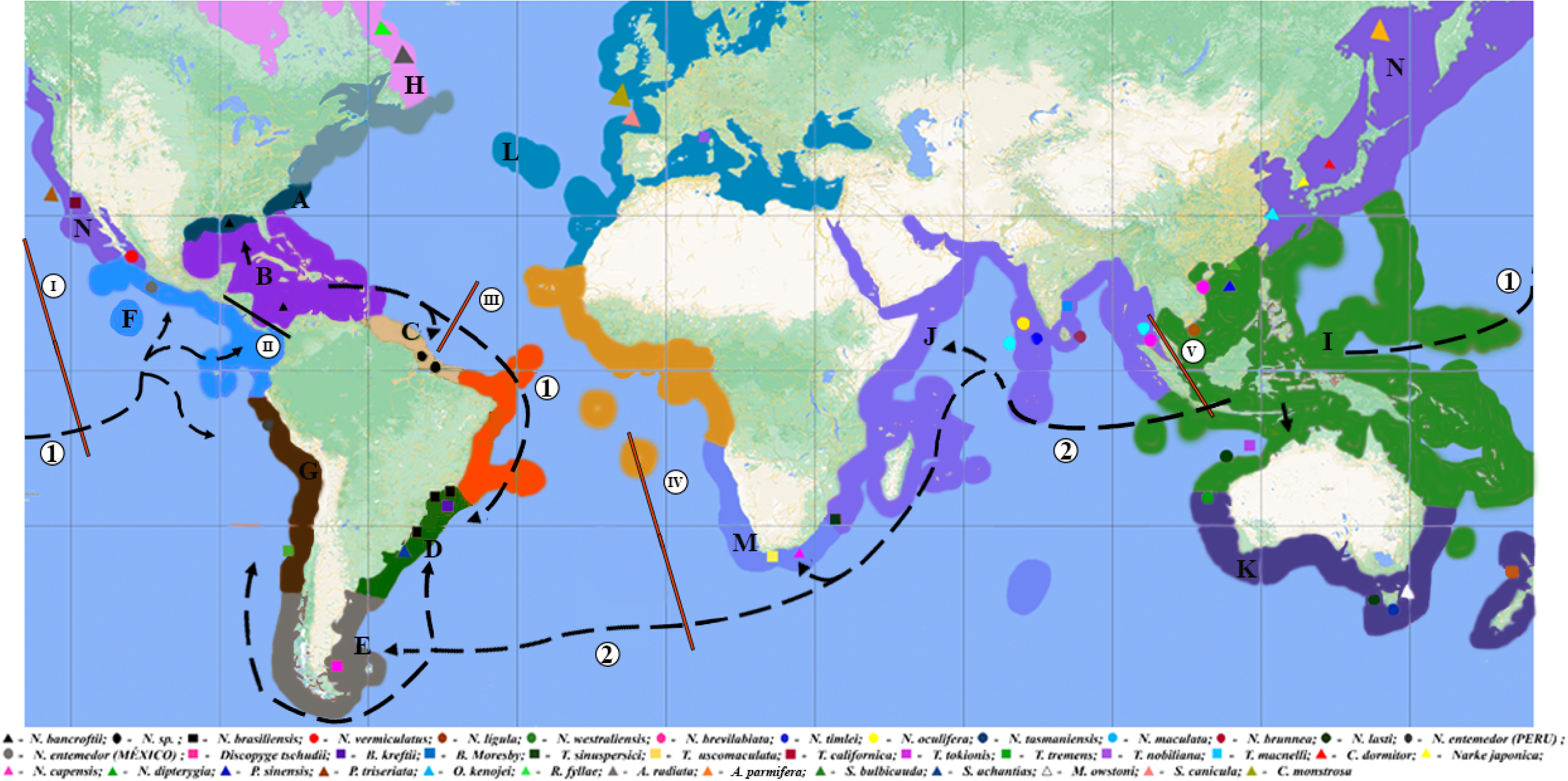
Distribution of the marine biogeographic realms and the species included in the present study. The letters and their respective colors correspond to the limits of the biogeographic realms and marine province defined by Spalding et al. (2007). I – East Pacific Barrier, II – Isthmus of Panama Barrier, III - Mid-Atlantic Barrier, IV – Indo-Pacific Barrier. 1 – Dispersal route of the narcinids from the Indo-Pacific to the Pacific through the East Pacific Barrier, and to the Caribbean prior to the closure of the Isthmus of Panama, 2 – Dispersal route of the narcinids from the Indo-Pacific to the Atlantic Ocean through the western Indo-Pacific and South Africa (see Table S1).

Additional narcinid and torpediniform sequences were obtained from GenBank and BOLD, together with sequences of other rays, sharks, and chimeras (forming the outgroup), and were included in the database for the present study, which had a total of 215 sequences for analysis (Table S1; Fig. 1). The collection of samples was authorized by the Biodiversity Authorization and Information System (SISBIO) through permanent license 12773-1e license 78993-3-1), and the Brazilian National System for the Management of Genetic Heritage and Associated Traditional Knowledge (SISGen: AD61D8E).

### Laboratory procedures, DNA extraction and Polymerase Chain Reaction (PCR)

The laboratory procedures, including the extraction of the DNA and the Polymerase Chain Reactions (PCRs), were conducted at the Genetics and Biotechnology Laboratory of the Capanema campus of the Federal Rural University of Amazonia. The amplicons were sequenced at the Bragança campus of the Federal University of Pará.

The total DNA was extracted using the Wizard Genomics DNA Purification kit (Madison, WI, USA) with the mousetailprotocol. Following the extraction and quantification of the DNA, the Polymerase Chain Reaction (PCR) was implemented using the primers described by Ward et al. (2005) for the mitochondrial Cytochrome Oxidase subunit I (COI) gene. The final reaction volume and the amplification conditions were the same as those used by Rodrigues-Filho et al. (2020).

The positive PCRs were purified using the Polyethylene Glycol 20% (PEG 20%) protocol, adapted from Dunn and Blattner (1987). The products were then precipitated for the sequencing reaction, which was mediated by the Big Dye kit (ABI PrismTM Dye Terminator Cycle Sequencing Reading Reaction – Applied Biosystems, Foster City, CA, USA), and then sequenced in an ABI 3500 Genetic Analyzer automatic sequencer.

The quality of the sequences obtained through this processing was verified in the Chromas software (available at: http://technelysium.com.au/wp/chromas/), and the sequences were edited and aligned in BioEdit 7.2.5 (Hall 1999), using the CLUSTALW automatic alignment tool (Thompson et al. 1994). The sequences were subsequently conferred visually to identify potential incongruities.

### Phylogenetic analysis

The present study was based on two alternative phylogenetic reconstruction approaches: Maximum Likelihood (ML), run on the IQ-TREE web server (Trifinopoulos et al. 2016; link: http://iqtree.cibiv.univie.ac.at/), and Bayesian Inference (BI), which was implemented in BEAST 2.7.5 (Bouckaert et al. 2019). Initially, the nucleotide substitution models for these two approaches were inferred in IQ-TREE (ML) and BEAST 2.7.5 (BI), using the bModelTest v1.1.2 package (Bouckaert and Drummond 2017). This process identified the TIM3e+I+G evolutionary model for the ML phylogenetic reconstruction and the TN93+I+G model for the Bayesian Inference.

In the ML analysis, branch support was determined using ultrafast bootstrap (UFBoot; Minh et al. 2013), the SH-aLRT test (Guindon et al. 2010), and the aBayes test (Anisimova et al. 2011), which were run on the IQ-TREE web server, with 1000 bootstrap replicates, following the program’s default setting. The confidence of the clades was inferred based on the minimum support for each procedure, i.e., SH-aLRT ≥ 80, aBayes ≥ 0.90, and UFBoot ≥ 95 (Guindon et al. 2010; Minh et al. 2013).

In the case of the Bayesian Inference, the Markov Chain Monte Carlo (MCMC) was run with one cold and three hot chains, to calculate the mean spacing of the trees, in order to weight each tree proportionally by its *a posteriori* probability (PP), based on a run of 60 million generations, which was sampled every 1000 generations, with the first 10% of the samples being discarded as burn-in. The quality of the run was assessed in Tracer v1.7 (Rambaut et al. 2018), to verify that the Effective Sample Size was at least 200 (ESS>200), with the result of the run then being submitted to TreeAnnotator (Rambaut and Drummond 2013) a package of the BEAST 2.7.5 program (Bouckaert et al. 2019), for the construction of the trees.

In addition to the phylogenetic inferences, matrices of genetic *p* distances were compiled in MEGA X (Kumar et al. 2018). Phylogenetic haplotype networks were compiled using the Neighbour-Net algorithm in SplitsTree 4.16.1 (Hudson and Bryant 2024) to permit the visualization of the evolutionary relationships of each species.

### Species delimitation

Three methods of species delimitation were applied in the present study to validate the evolutionary lineages of the Narcinidae defined in the analyses. The first method was the Automatic Barcoding Gap Discovery (ABGD) tool of Puillandre et al. (2012). In the case of this approach, six distinct runs combined different distance matrices (K80, JC, and simple distance) with two relative gaps (X = 1.0 and 1.5). In this approach, only the recursive partitions were evaluated, given that they permit different gap thresholds between the taxa, with two partitions being selected for comparison here – (i) the recursive partition with the smallest possible number of detected species (RP 1), and (ii) the recursive partition with the largest possible “adequate” number of species (RP 2). As the ABGD may produce an ample range of delimitations, due to the variation in the prior maximal intraspecific distance, P (Puillandre et al. 2012), the RP 2 was selected based on the pre-established species definitions (Supplementary Data 1), and the other species delimitation and phylogenetic analyses. All the other parameters were set at the default value provided by the program (Puillandre et al. 2012; link: https://bioinfo.mnhn.fr/abi/public/abgd/abgdweb.html).

The Poisson Tree Processes (PTP) method, based on the Bayesian model (bPTP) of species delimitation (Zhang et al. 2013), was run on the https://species.h-its.org/web server (Pons et al., 2006; Fontaneto et al., 2007; Monaghan et al. 2009). This procedure models the speciation rate directly from the number of substitutions, without needing an ultrametric tree, with the species being delimited based on the phylogenetic species concept (Zhanget al. 2013). The input tree (ML) for the bPTP was generated by IQ-TREE. The MCMC run had 500,000 generations, with a burn-in of 10%, with all the other parameters being set at the default values provided by the program. This analysis recovered two distinct trees, one with the support derived from the values of *a posteriori* probability (bPTP-BI) and the other, with the Maximum Likelihood values (PTP-ML).

In the final approach, a Neighbor-Joining (NJ) tree was obtained from MEGA X (Kumar et al. 2018) as a phylogenetic tool for the comparison of the different methods of species delimitation. The nucleotide substitution model used in this approach was established by the Find Best DNA/Protein Models, run in MEGA X (GTR+G+I), while the robustness of the clades was evaluated based on 1000 bootstrap replicates (Felsenstein 1985).

### Divergence times estimative and biogeographic reconstruction

The estimates of the Time of Most Recent Common Ancestor (TMRCA) between the members of the family Narcinidae were obtained in BEAST 2.7.5 (Bouckaert et al. 2019), using the substitution model (TN93+I+G) established by the bModelTest v1.1.2 package (Bouckaert and Drummond 2017). The Calibrated Yule Model was used as the tree prior, with a Strict Clock (Drummond et al. 2006). All the other parameters were set previously to the program’s default values. The following calibration points (see Aschliman et al. 2012) were used for this analysis – (i) approximately 213.4 Million years ago (Mya) for the separation of the sharks and rays, (ii) approximately 187.8 Mya for the divergence of the skates from the other rays, and (iii) approximately 164.2 Mya for the separation of the electric rays from the thornbacks. The Markov Chain Monte Carlo (MCMC) was based on a run of 60 million generations, which were sampled every 1000 generations, with 10% of the samples being discarded as burn-in. The results of this analysis were inspected in Tracer v1.7 (Rambaut et al., 2018) to determine the Effective Sample Size (ESS>200). The consensus tree derived from the BEAST run was summarized in the TreeAnnotator package (Bouckaert et al. 2019) to allow for the inclusion of the calibration times of the tree branches.

The biogeographic history, dispersal routes, and probable ancestral area of the Narcinidae and the other associated species (outgroup), were reconstructed using the log files and consensus tree of the divergence times, using the Bayesian Binary MCMC (BBM) method for the reconstruction of the ancestral state, which was run in RASP 4.2 (Yu et al. 2020). This method uses the trees of the *a posterior* distribution of the Bayesian analysis as the input for the inference of ancestral distributions through a complete Bayesian hierarchical approach. The MCMC analysis of the BBM was run with 10 chains, which were run simultaneously for 50 million generations, with a sampling frequency of once every 100 generations and a burn-in of 10%. The fixed Jukes-Cantor model was used for BBM analysis, given that it was the default procedure in the program. This reconstruction was based on the biogeographic realms and marine provinces delimited by Spalding et al. (2007) for the definition of the areas occupied by the study species (Table S1; Fig. 1), their probable ancestral areas, and their likely routes of dispersal and speciation.

## RESULTS

### Phylogenetic inferences and genetic distance

The results of the Bayesian analyses indicate that the narcinid species investigated in the present study were distributed in four groups, well-supported clades, with high *a posteriori* probabilities, which were denominated: (i) the American *Narcine* clade (composed by Pacific and Atlantic Lineages), (ii) the Central/Western Indo-Pacific *Narcine* clade, (iii) the *Narcinops* clade, and (iv) the *Bentobatis/Discopyge* clade (Fig. 2A and Fig. 3). In the phylogenetic relationships recovered within the American clade, the Atlantic species are phylogenetically closest to *Narcine entemedor* Jordan and Starks, 1895, which is the sister group of the Atlantic American lineages (PP = 1.0). This analysis also revealed a clear dichotomy between the *N. entermedor* individuals from Peru and Mexico (PP = 1.0). *Narcine vermiculatus* Breder, 1928 was also recovered as the most divergent lineage of the species that make up the American clade (Fig. 2A).

**Fig. 2.**
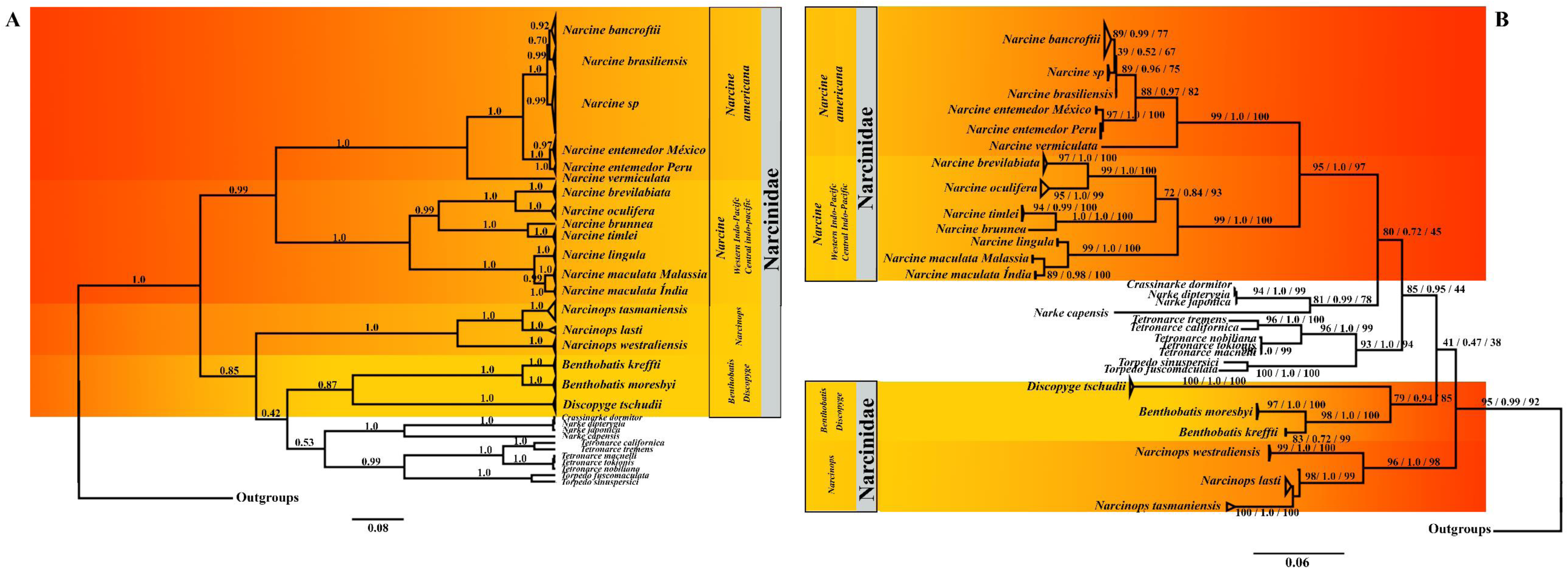
Phylogenetic relationships within the family Narcinidae, based on the (A) BI and (B) ML analyses. The Posterior Probabilities (PP) are shown above the branches of each clade in the BI tree. The SH-aLRT, aBayes, and UFBoot values are shown in the ML tree above the branches of each clade, in this order (SH-aLRT/aBayes/UFBoot).

**Fig. 3.**
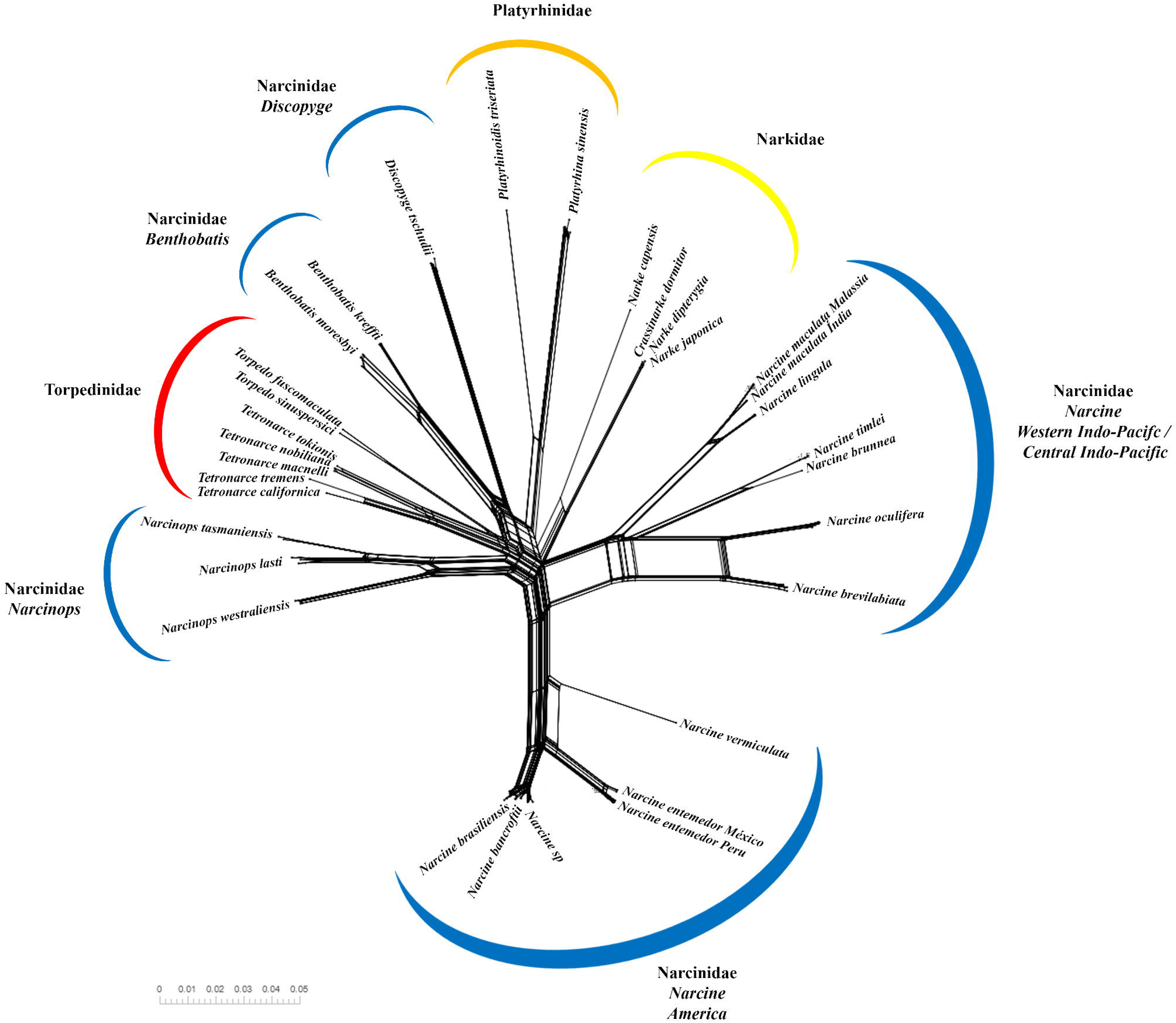
Analysis of the Neighbor-Net phylogenetic network. The scale bar represents the number of nucleotide substitutions per site.

In the American Atlantic, the *Narcine sp.* clade was clearly separated from the *Narcine brasiliensis*/*Narcine bancroftii* clade (PP = 1.0), and thus appeared to be the most divergent of the three species (Fig. 2A). Two monophyletic subclades were also observed in each species of the *N. brasiliensis*/*N. bancroftii* clade (PP: 0.70), which were well defined in both the *N. brasiliensis* (PP = 0.99) and the *N. bancroftii* lineages (PP = 0.92).

In the Central/Western Indo-Pacific clade, *Narcine brevilabiata* Bessednov, 1966, *Narcine oculifera* Carvalho, Compagno and Mee, 2002, *Narcine timlei* (Bloch and Schneider, 1801), and *Narcine brunnea* Annandale, 1909 were recovered as the most closely-related species, with *Narcine lingula* Richardson, 1846 and *Narcine maculata* (Shaw, 1804) as the sister group of these species (PP = 1.0). In the case of *N. maculata*, a clear dichotomy was identified between the individuals from India and Malaysia (PP = 0.99).

In the case of the *Narcinops* clade, as in the *Bentobatis/Discopyge* clade, the phylogenetic inferences recovered a well-supported arrangement, indicating that this was the most ancestral narcinid group, which formed a clade with the species of the families Narkidae and Torpedinidae (Fig. 2A). All the phylogenetic reconstructions recovered in the present study also indicated the possible separation of the *Narcine* lineages from the American and Central/Western Indo-Pacific regions (PP = 0.99), as well as the paraphyly of the family Narcinidae, observed in the phylogenetic proximity between the other genera of this family (*Narcinops*, *Bentobatis*, and *Discopyge*) and representatives of the families Narkidae and Toperdinidae (Fig. 2A).

The Maximum Likelihood (ML) tree also recovered the four clades identified in the Bayesian tree (Fig. 2B), with high levels of support for each clade (SH-aLRT > 90, aBayes > 0.90, UFBoot > 90), as well as the possible separation of the American and Central/Western Indo-Pacific *Narcine* lineages (SH-aLRT = 95, aBayes = 1.0, UFBoot = 97). The ML tree also placed the narkid clade as the sister clade of *Narcine*, while the Torperdinidae, *Benthobatis, Discopyge*, and *Narcinops* clades were the most external in the tree (Fig. 2B). Even so, it is important to note that the topology of the ML tree does not confirm the paraphyly of the family Narcinidae, given the low levels of statistical support observed in the most basal nodes of this tree (Fig. 2B).

### Neighbour-Net phylogenetic network

The Neighbor-Net analysis produced a well-structured network of phylogenetic relationships, which also defined the clades observed in the phylogenetic trees, i.e., American *Narcine*, Central/Western Indo-Pacific *Narcine*, *Narcinops*, and *Benthobatis/Discopyge* (Fig. 3). The phylogenetic relationships observed in this analysis are consistent overall, and broadly similar when comparing the ML and BI phylogenetic trees. In particular, the clear separation of the American and Central/Western Indo-Pacific *Narcine* lineages, as well as the presence of the dichotomies in *N. entemedor* and *N. malculata* (Fig. 3). The paraphyly of the Narcinidae is also apparent, given the closer phylogenetic relationships between the genera *Benthobatis* and *Discopyge*, and the representatives of the family Torpedinidae.

### Genetic distance

The analysis of genetic distance also confirms the existence of four narcinid clades (Table 1 and Table S2), separated by mean distances of approximately 17.2% (American *Narcine* x *Narcinops*), 18.2% (American *Narcine* clade x Central/Western Indo-Pacific *Narcine* clade), and 19.1 % (Central/Western Indo-Pacific x *Narcinops*). These distance values are very close to those found in the comparisons between the families of the order Torpediniformes and other rays (Table S2).

**Table 1.** Matrix of the *p* distances (%) between the sequences of the narcinid species, based on their biogeographic regions. The mean genetic distances found between the species of the American *Narcine* clade are shown in light gray, with the *Narcine* clade of the Central/Western Indo-Pacific in dark gray, the *Narcinops* clade in light yellow, and the *Bentobatis*/*Discopyge* clade in dark yellow. The mean values for the *Narcine* species found in the Atlantic are shown in bold script.

Relatively higher distance values were recorded between the *Bentobatis*/*Discopyge* clade and the *Narcine* clades than in the comparisons between the clades (American *Narcine* x *Bentobatis* – 19.0 %; American *Narcine* x *Discopyge* – 20.8%; Central/Western Indo-Pacific *Narcine* x *Bentobatis* – 20.1%; Central/Western Indo-Pacific *Narcine* x *Discopyge* – 21.1%). High values were also observed between the *Narcine* species from the American and Central/Western Indo-Pacific regions, which were compatible with those recorded in the comparisons with families of other orders (Table 1 and Table S2). In fact, the genetic distances recorded between the *Narcine* species of the Central/Western Indo-Pacific ranged from 1.9% to 20%. These values were well above those observed in the *Narcine* species of the American clade (i.e., *N. sp.* x *N. brasiliensis* x *N. bancroftii* = 0.7–1.1 %).

It is important to note here that *N. entemedor* (from the Pacific) presented a high level of differentiation from the Atlantic *Narcine* lineages (4.8–5.6 %), although they were well below the distances observed in the other *Narcine* species of the Central/Western Indo-Pacific clade and in the *Narcinops* clade. By contrast, the distance between *N. entermedor* and *N. vermiculatus* was 9.5%. The dichotomy between the *N. entemedor* individuals from Peru and Mexico had a genetic distance (0.9%) that was like that found between the species of the Atlantic, whereas the dichotomy observed in *N. maculata* was more pronounced, with a distance of 1.9% (Table 1).

### Species delimitation

The species delimitation analysis of the family Narcinidae revealed clusters of highly similar candidate species, with very little variation, due to the mixing and division of some species (Table S3). In the case of the ABGD, the K80 (X=1.0 and 1.5) and JC (X=1.0 and 1.5) combinations resulted in identical recursive partitions, with the number of clusters of candidate species ranging from 15 to 20. The recursive partition with the smallest number of clusters (RP1; P=0.007743) recovered 16 species clusters, in a pattern similar to that established in Table 1, except for *Narcine* sp., *N. brasiliensis*, and *N. bancroftii*, which were included in a single cluster, and *N. entemedor*, in which the individuals from Mexico and Peru were separated into different clusters. For the recursive partitions with the largest number of clusters (RP2; P=0.004642; 18 clusters), all the species were recovered in distinct clusters, except for *Narcine* sp., *N. brasiliensis*, and *N. bancroftii*, included in a single cluster, and *N. entemedor* and *N. maculata*, which were each separated into two clusters (Table S3). The ABGD analysis for the combinations with the Simple Distance (SD) model varied little in the number of clusters (SD: X=1.0 – 14–15 clusters and X=1.5 – 13–15 clusters). All the recursive partitions recovered for the SD (X=1.0; P=0.035938) had the same number of clusters (14). A similar result was obtained for the SD-combination (X=1.5; P=0.035938), but with 13 clusters (Table S3).

The bPTP-ML and bPTP-BI analyses returned high support values for the majority of the clusters, which decreased as the individuals of a species recovered in distinct clusters. The bPTP-ML recovered17 species clusters, which mixed *Narcine* sp., *N. brasiliensis*, and *N. bancroftii*, while *N. lingula* was separated into three distinct clusters. The bPTP-BI recovered the largest number of clusters (26) recorded in any of the species delimitation analyses. These 26 clusters were derived from the separation into distinct clusters of individuals of *N. brevilabiata* (two clusters), *N. lingula* (three clusters), *N. maculata* (three clusters), *N. westraliensis* (three clusters), and *D. tschudii* (five clusters). The bPTP-BI also recovered a mixture of individuals of *Narcine* sp., *N. brasiliensis*, and *N. bancroftii*. For comparison, the NJ tree recovered 19 monophyletic clusters (Bs > 80), which separated individuals of *N. entemedor* and *N. maculata* (each with two clusters), as well as *Narcine* sp., *N. brasiliensis*, and *N. bancroftii* (Table S3).

### Biogeographic reconstruction and divergence times estimates

The results of the biogeographic reconstruction, together with the divergence time estimates indicate that the probable ancestral area of the Torpediniformes was the Central Indo-Pacific realm (96.45%), from which the various lineages of the genus *Narcine* dispersed, including the clades from the Americas and the Indo-Pacific. The *Narcinops/Bentobatis/Discopyge*/Narkidae/Torpedinidae clade dispersed from this region at around 153 Mya (node I; Fig. 4; Table 2).

**Fig. 4.**
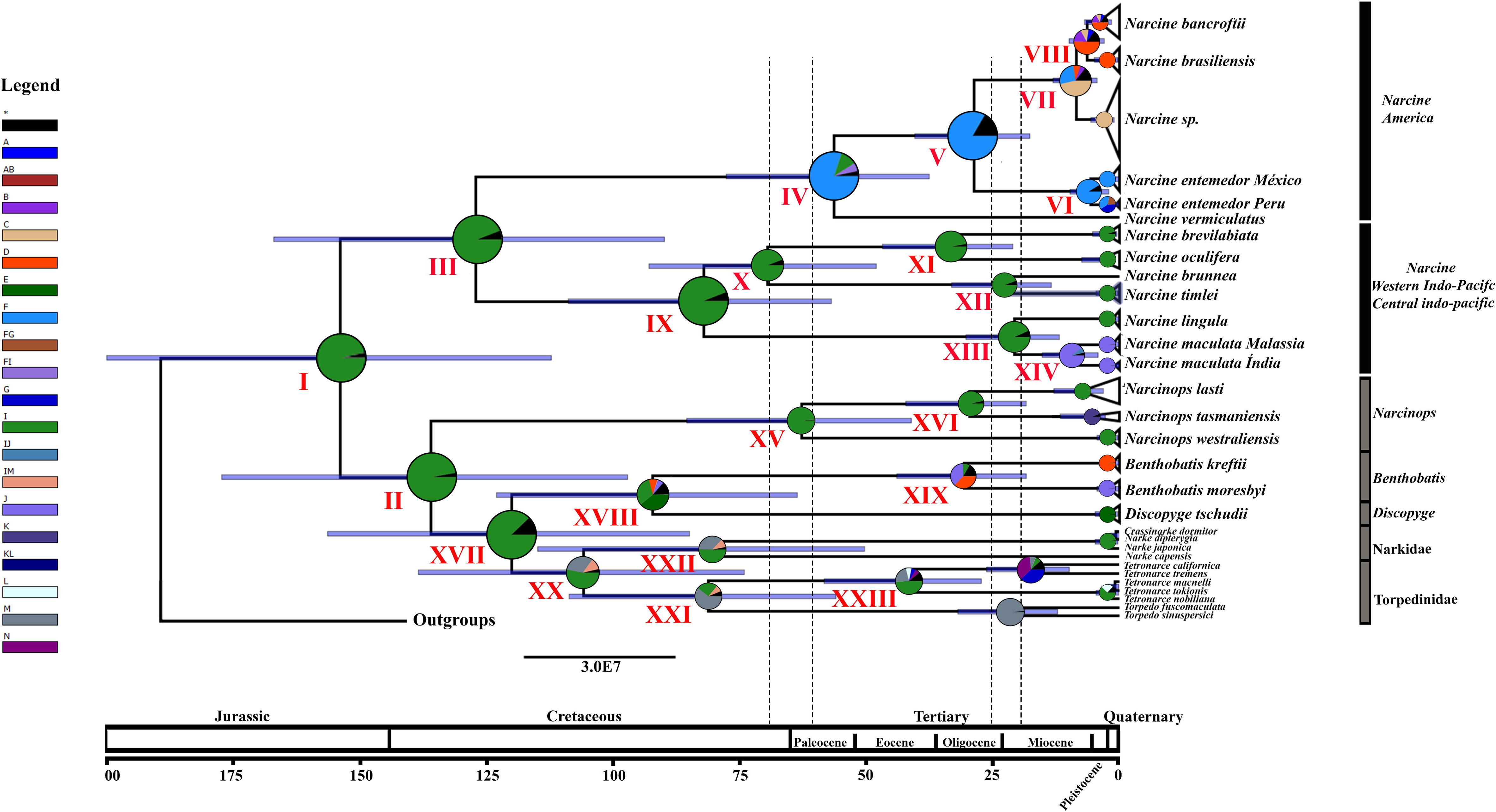
Estimate of the TMRCA and reconstruction of the biogeographic history of the family Narcinidae based on the BBM method, run in RASP 4.2 (Yuet al., 2020). The circles at each node of the tree represent the probability of the possible ancestral area, which is cited in the caption to the left of the tree (see Table 2, Table S1). The vertical bars to the right indicate the genetic groups identified here. The estimates of divergence times are shown on the horizontal scale bar located below the tree.

**Table 2.** Estimated Divergence Times (DT (HPD)) for the principal narcinid lineages, with the probable dispersal routes and the number of dispersals, vicariant, and extinction events identified in the separation of some lineages. (->) – dispersalbetween realms. () - dispersal within the same realm. (|) – separation event. A - Warm temperate Northwest Atlantic; B - Tropical Northwestern Atlantic; C - North Brazil Shelf; D - Warm temperate Southwestern Atlantic; E - Magellanic; F – Warm temperate Southwestern Atlantic; G - Warm temperate Southeastern Pacific; I - Central Indo-Pacific; J - Western Indo-Pacific; K - Temperate Australasia; L - Temperate Northern Atlantic; M - Temperate Southern Africa; N - Temperate Northern Pacific. BBM – Bayesian Binary MCMC; Disp. – Dispersal; Vic. – Vicariance; Ext. – Extinction.

The ancestors of the genus *Narcinops* and the *Bentobatis/Discopyge*/Narkidae/Torpedinidae clade emerged at ∼135.95 Mya (node II; Fig. 4; Table 2), also in the Central Indo-Pacific (97.25%). *Narcine* emerged and began to diversify at ∼126.96 Mya, with at least one vicariant and dispersal event that separated two ancestral lineages, which are now distributed in the American and Central/Western Indo-Pacific realms (node III; Fig. 4; Table 2). The probability of the possible dispersal route of the group (0.7052) indicates that the ancestral lineage separated in and dispersed from the Central Indo-Pacific realm. Following this separation, one likely dispersal event began in the Central Indo-Pacific realm and moved eastward to the tropical East Pacific realm, while a second event occurred between the Central and the Western Indo-Pacific realms (nodes IV–XIV; Fig. 4; Table 2).

The analyses presented here recovered at least one vicariant event for each dispersal route, which separated the American narcinids into two distinct lineages at ∼56.23 Mya (node IV; Fig. 4; Table 2). The lineages of the Central/Western Indo-Pacific, in turn, diversified into the Central Indo-Pacific realm (*N. lingula/N. maculata*) and Western Indo-Pacific lineages (*N. brevilabiata*/*N. oculifera/N. brunnea/N. timlei*) at ∼81.89 Mya (node IX; Fig. 4; Table 2). The ancestral *Narcinops*, *Bentobatis*, and *Discopyge* remained in a distinct clade (distant from the *Narcine* species of the American and Central/Western Indo-Pacific realms), which were more closely-related to the representatives of the families Narkidae and Torpedinidae, which emerged at ∼135.95 Mya (node XV; Fig. 4; Table 2).

### Diversification of the American narcinids

The ancestral lineage of the American narcinids dispersed from the Tropical East Pacific realm toward the tropical Atlantic realm (node IV; Fig. 4; Table 2). Prior to its arrival in the Tropical Atlantic, a vicariant event occurred at ∼56.23 Mya, which separated the lineages that gave rise to *N. vermiculatus* and the ancestral lineage from the *N. entemedor* complex and the Atlantic *Narcine*, which emerged at ∼28.54 Mya. This diversification was driven by processes of dispersal and vicariance that occurred within the tropical East Pacific and concluded with the dispersal of the *Narcine* lineages to the tropical Atlantic realm (nodes IV–V; Fig. 4; Table 2).

Two lineages were identified in the *N. entemedor* complex, which were derived from a divergence process, based on vicariance and dispersal, which began around ∼4.19 Mya (node VI) off the Pacific coast of Mexico (Tropical East Pacific), from where the first lineage was identified. This lineage dispersed subsequently to the warm temperate southeastern Pacific, off the coast of Peru, where the second lineage settled.

The probable dispersal route of the ancestral lineage of the Atlantic *Narcine* species was from the Warm Temperate Southwestern Atlantic toward the province of the North Brazil Shelf, where the *Narcine* sp. lineage began to diverge from the ancestor of *N. brasiliensis*/*N. bancroftti* at ∼8.43 Mya (node VII), with the latter lineage then dispersing from the North Brazil Shelf province to ward the Warm Temperate Southwestern Atlantic. Following this event, the ancestor of *N. bancroftti* and *N. brasiliensis,* separated at ∼6.29 Mya (node VIII), by dispersing from the Warm Temperate Southwestern Atlantic province (*N. brasiliensis*) toward the Tropical Northwestern Atlantic province, i.e., *N. bancroftti* (Fig. 4; Table 2).

### Diversification of the *Narcine* species from the Central/Western Indo-Pacific realm

The ancestor of the genus *Narcine*, which inhabited the Central/Western Indo-Pacific realm fragmented into the lineages that became *N. timlei*, *N. oculifera*, *N. maculata*, *N. brunnea*, and *N. brevilabiata* at ∼ 81.89 Mya, from a center of dispersal in the Central Indo-Pacific realm. This diversification involved events of both dispersal and vicariance, following the route from the Central Indo-Pacific to the Western Indo-Pacific realm (node IX; Fig. 4; Table 2).

*Narcine oculifera* diverged from *N. brevilabiata* at around 33.26 Mya, through the separation of the Central Indo-Pacific species (*N. brevilabiata*) from that (*N. oculifera*) of the Western Indo-Pacific (node XI). A similar process separated *N. lingula* from the Central Indo-Pacific, from *N. maculata* (western Indo-Pacific) at ∼20.69 Mya (node XIII), while *N. brunnea* and *N. timlei* diverged at ∼22.83 Mya in the western Indo-Pacific realm (node XII). The different *N. maculata* lineages observed in India and Malaysia diverged at ∼9.37 Mya in the Western Indo-Pacific (node IX; Fig. 4; Table 2).

### Diversification of the species of the genera *Narcinops*, *Benthobatis*, and *Discopyge*

The ancestor of the *Narcinops*/*Bentobatis/Discopyge*/Narkidae/Torpedinidae clade emerged in the realm of the Central Indo-Pacific, with the *Narcinops* lineage separating from the ancestral of *Bentobatis/Discopyge*/Narkidae/Torpedinidae lineage 135.95 Mya. This was probably followed by dispersal to the Australia realm, where *Narcinops westraliensis* (McKay, 1966), *Narcine lasti*, and *Narcine tasmaniensis* (Richardson, 1841) emerged through successive vicariant and dispersal events (nodes XV–XVII). This sequence of events began at around 62.64 Mya, with the separation, still in the Central Indo-Pacific realm, of the most divergent species of this clade (*N. westraliensis*) from the ancestor of *N. lasti/N. tasmaniensis* (node XVI), with subsequent dispersal to the region more to the south of Australia and vicariance of the lineage that gave rise to *N. tasmaniensis* at ∼28.1 Mya(node XVII; Fig. 4; Table 2).

The dispersal route of *Benthobatis* and *Discopyge* to the American continent was different from that established for the American *Narcine*. This dispersal event began with the diversification of the Narkidae/Torpedinidae and *Benthobatis/Discopyge* lineages at ∼119.87 Mya (node XVIII), which originated in the Central Indo-Pacific realm and moved toward temperate southern Africa, reaching the American coast in the Magellanic realm. The ancestor of *Benthobatis/Discopyge* diverged into two genera at ∼92.01 Mya (node XIX), dispersing in opposite directions, toward the Magellanic realm (*Benthobatis kreffti* Rincón, Stehmann and Vooren, 2001*/Discopyge tschudii* Heckel, 1846) and the Western Indo-Pacific (*Benthobatis moresbyi* Alcock, 1898). *Benthobatis kreffti* dispersed to the warm temperate southwestern Atlantic, whereas *B. moresbyi* settled in the realm of the western Indo-Pacific, with *D. tschudii* establishing its distribution in the Magellanic (node XIX-XX; Fig. 4; Table 2).

The separation of the lineage of the family Torpedinidae from Narkidae was estimated to have occurred at approximately 105.82 Mya (node XX), with dispersal toward to the realm of temperate Southern Africa. The same route was observed during the separation of the genera *Torpedo* and *Tetronarce* at ∼81.06 Mya (node XXI). The separation of the lineages of the *Tetronarce* species found in the Magellanic and the Central Indo-Pacific realms was estimated to have occurred at ∼41.98 Mya (node XXIII; Fig. 4; Table 2).

## DISCUSSION

### Biogeographic History of the Narcinidae

The TMRCA inferences derived from the data obtained in the present study provided dating estimates that are consistent with those of previous studies, which concluded that the Torpediniformes emerged during the Cretaceous period (Aschliman et al. 2012: 164.2 Mya, Height posterior density (HPD): 150.2–179.7 Mya; Villalobos-Segura and Underwood 2020: 106.04 Mya, HPD: 86.64–126.61 Mya). During this period, most of the planet’s shallow continental waters were associated with the Tethys Sea, an enormous waterway that stretched from the Central Atlantic to the region now known as the western Pacific (Aschliman et al. 2012). The vast majority of the modern batoids occupy this type of shallow-water environment, with the dispersal to deeper waters being limited by the eventual competition with the skates (Siverson and Cappetta 2001; McEachran and Aschliman 2004).

The most recent common ancestor of the electric rays was dated to ∼153.80 Mya (HPD: 112.03–199.89 Mya), which corresponds to the onset of the Cretaceous, a period marked by intense tectonic plate movements and substantial global increases in temperatures, driven by the reduction of the polar ice caps, together with the widening of the bottom of the Atlantic Ocean, which resulted in an abrupt increase in sea levels during the upper Cretaceous (Miller et al. 2003; Gale et al. 2020). As inferred by a number of previous studies (Guinot et al. 2012; Gales et al. 2024), the period between the end of the Cretaceous and early Paleocene (K/Pg), ∼65.50 Mya, was marked by rapid radiations in the taxa that were able to persist through this period of extinction. The oldest fossil attributed to an electric ray (Torpediformes) represents the only batoid family to appear during the Danian Age (∼66–61.6 Mya), whereas other extant families only arose later in the Paleocene (Aschliman et al. 2012; Cappetta 2012). The TMRCA identified in the present study indicates that this group of rays radiated rapidly, given the long internal branches, followed by multiple ramifications of species in the more external branches of the tree (Fig. 2 and Fig. 4; see Aschliman et al. 2012). This initiated the rapid diversification of the principal lineages of the families Narcinidae, Narkidae, and Torpedinidae between the late Cretaceous and early Paleocene (Fig. 4), which is consistent with the evidence that the radiation of all the modern clades of batoids occurred in the lower-middle Paleogene (Aschliman et al. 2012; Villalobos-Segura and Underwood 2020).

Several studies have highlighted the importance of processes such as vicariance and the dispersal of lineages for establishment of the present-day diversity and distribution of aquatic organisms (Floeter et al. 2008; Cowman and Bellwood 2013; Jurado-Riveira et al. 2017). The hypotheses proposed to account for these patterns include: 1) the theory of the origin of biodiversity through the Tethys Sea, 2) the colonization of the Pacific through the East Pacific Barrier and the Caribbean prior to the closure of the Isthmus of Panama, as well as 3) the colonization of the Atlantic from the Indian Ocean via South Africa (Rocha 2003; Rocha et al. 2007; Pellissier et al. 2014).

These theories point to the probable periods of diversification and dispersal of the ancestral lineages of the oceans’ present-day fauna. For example, the theory of the relict fauna of the Tethys Sea predicts that the ancestral lineages of the American continent were derived from the Central Indo-Pacific realm between 12 and 18 Mya. By contrast, the colonization of the Atlantic through the opening of the isthmus has been dated to around 3.1 Mya (Coates and Obando 1996), reflecting the greater phylogenetic proximity (sister taxa) of the fauna from the central Atlantic and eastern Pacific (Sales et al. 2019; Lima et al. 2020).

The region known as the Coral Triangle, in the Central Indo-Pacific realm, is considered to be an important biodiversity hotspot, which has remained stable for millions of years, supporting the origin of an enormous array of Indo-Pacific species (60% of the region’s reef-dwelling fauna), which have subsequently dispersed across the world’s oceans, including the ancestor of the modern electric rays (Carpenter et al. 2010; Barber et al. 2011). This association between areas of high species richness (Central Indo-Pacific) and the origin and dispersal of lineages has been well documented in both vertebrates (Summerer et al. 2001; Lessios and Robertson 2006; Floeter et al. 2008; Barber et al. 2011; Cowman and Bellwood 2013; Tornabene et al. 2015) and invertebrates (Ulloa et al. 2017). The narcinids represent one of the new taxonomic groups to emerge in this region (node I–III, Figure 4), together with other elasmobranch species (Daly-Engel et al. 2012; Sales et al. 2019), although there are some differences in relation to the exact center of origin and diversification (Gales et al. 2024).

The separation of the narcinid lineages from the Central Indo-Pacific and American realms occurred at ∼126.96 Mya (HPD: 89.70–166.90 Mya). In the Western Indo-Pacific realm, the separation of *N. lingula* and *N. maculata* dates to ∼20.69 Mya (HPD: 11.84–30.21 Mya), which is close to the estimated origin of the Indo-Pacific barrier, that began to form in the middle Miocene (16–8 Mya) as a result of the collision of the Australian and Eurasian plates. This process resulted in a reduction in the flow of water in the region between the present-day Indian and Pacific oceans. This barrier was influenced profoundly by the glaciations of the Pleistocene, when the sea level reached approximately 130 m below that of the present day (Bowen et al. 2016). This same hypothesis has been used to account for the separation of *N. timlei* and *N. brunnea,* which occurred at ∼22.83 Mya (HPD: 13.41–33.09 Mya) and *N. oculifera* and *N. brevilabiata*, at ∼33.26 Mya (HPD: 21.02–62.79 Mya).

The lineages that originated the *Narcinops* species began to diverge from the *Bentobatis/Discopyge*/Narkidae/Torpedinidae clade lineage at around 135.95 Mya, still in the ancestral area of the Central Indo-Pacific realm. Previous studies indicated the presence of barriers between the Indo/Australian archipelago and the Indian and Indo-Pacific oceans, which provoked vicariant events between these regions that extended up until the Miocene (approximately 15 Mya). The glaciations that occurred during the transition from the Pleistocene to the Holocene also resulted in alterations in sea level, which had a direct influence on the structuring of the populations of the species of the Indo/Australian archipelago and Central Pacific (Cowman and Bellwood 2013). However, the diversification events observed in the species of this area are more ancient (between 28 and 60 Mya). Cowman and Bellwood (2013) identified later vicariant events in this region, which they linked to events on a global scale, influenced by climate change or shifts in ocean currents. The present study found evidence that the emergence and diversification of the modern electric ray taxa occurred at 25– 20 Mya, which is consistent with the emergence of barriers and the diversification events mentioned above for the Central/Western Indo-Pacific realm.

The separation of the lineages of the Central Indo-Pacific realm from those of the Tropical Eastern Pacific realm may have been associated with the Eastern Pacific Barrier, which limits the distribution of the tropical species (Gaither et al. 2016; Floeter et al. 2008). Even so, some taxa are able to maintain population connectivity across this barrier, as indicated by their low or non-significant Φ_ST_ values (Bowen et al. 2016).

Duncan et al. (2006) obtained findings closely like those of the present study in their global scale analysis of the shark *Sphyrna lewini*, based on genetic distances and haplotype networks. This study also identified the Cantreal Indo-Pacific realm as the ancestral area of the species, with dispersal events occurring during the Pleistocene in the direction of the central (Hawai’i) and eastern Pacific (Central America). The genetic discontinuity observed in *S. lewini* was also associated primarily with oceanic barriers (e.g., the Indo-Pacific barrier and the Isthmus of Panama). Similar evidence on the influence of barriers, including seasonal patterns, can be found in previous phylogeographic studies of sister groups, such as the species of the fish genera *Siganus*, *Chaetodon*, *Scarus* and *Halichoeres* (Bowen et al. 2016).

The separation of the *Narcine* lineages of the Atlantic from *N. entemedor* at ∼28.54 Mya (HPD: 17.59–40.30 Mya) would have been well before the closure of the Isthmus of Panama. Despite the controversies over the exact dating of this event, more recent studies indicate that the isthmus began to form between 38 and 28 Mya, and that the southern tip of Central America began to collide with South America during the upper Oligocene, with the isthmus closing completely by around 14–15 Mya (Bacon et al. 2015; Montes et al. 2015). In this case, the possible response of the *Narcine* lineages to enable the colonization of the Atlantic may have been associated with a discreet increase in sea level, of approximately 10 m, at around 24 Mya (Miller et al. 2013), which would have facilitated the colonization of the shallower areas of the continental shelf. This type of dispersal prior to the closure of the Isthmus of Panama has been recorded in other marine organisms and was responsible for the dispersal of the *Lolliguncula* lineages through the Atlantic Ocean (Costa et al. 2021) during the same period.

Probably about the same time when the ancestral of *Narcine* may have migrated to the region of the Atlantic through the opening in the isthmus, to become isolated completely from *N. entemedor* (Knowlton and Weigt, 1998; Bacon et al. 2015). This vicariant event likely also contributed to the separation of other species of marine organisms (Amorim and Costa 2018; García-de-León et al. 2018; Lima et al. 2020), including the elasmobranchs (Schultz et al. 2008; Chabot and Allen 2009; Sales et al. 2019).

Previous studies have already highlighted the morphological proximity between the species and lineages of the Western Atlantic and Central Pacific (Knowlton and Weigt, 1998; Bernardi and Lape 2005; Floeter et al. 2008). Floeter et al. (2008) demonstrated that 69% of the fishes of the Caribbean are phylogenetically closer to those of the Tropical Eastern Pacific than the Eastern Atlantic, which would be consistent with the direction of the dispersal of the ancestral lineage of *Narcine* from the Central Indo-Pacific realm to the Tropical Eastern Pacific realm and, ultimately, the American Atlantic, through the opening of the Isthmus of Panama.

It is important to note here that the divergence time between *N. entemedor* and the *Narcine* species of the American Atlantic, is still prior to the date proposed by Montes et al. (2015) for the complete closure of the Isthmus of Panama, even when the confidence interval is considered. This supports the conclusion that the separation of these lineages may have been more closely related to ecological and oceanographic factors, as proposed for some other marine organisms distributed between the tropical Atlantic and Pacific oceans. In their study of the genus *Octopus*, for example, Lima et al. (2020) found that dispersal and vicariance events between the Pacific and American Atlantic lineages occurred “during the process of the formation of the isthmus”, with the study species diverging at 5–18 Mya, when there was sufficient connectivity between the Caribbean and Pacific to maintain a dispersal route between these areas for some species up until the lower Pliocene.

The formation of the Isthmus of Panama resulted in a series of gradual changes that caused major shifts in oceanographic conditions (e.g., temperature, salinity, circulation, and productivity), which also affected the global climate. This likely influenced the genetic divergence of the various invertebrates through the formation of the physical and environmental barriers that arose in association with the formation of the isthmus, rather than its complete closure. A range of other marine organisms, such as the echinoids, crustaceans, fish, and molluscs (Marko 2002; Lessios 2008; O’Dea et al. 2016), in addition to elasmobranchs (Poortvliet al. 2015; Sales et al. 2019), were also affected by this process.

In the case of the *Narcine* species from the Atlantic, *Narcine* sp. initially separated from the ancestor of the other *Narcine* species (∼8.43 Mya) in the Tropical Atlantic realm, and then dispersed to the North Brazil Shelf province. At around 6.29 Mya, *N. bancroftii* diverged from *N. brasiliensis*, with *N. brasiliensis* dispersing to its current range in the Warm Temperate Southwestern Atlantic realm and *N. bancroftii* to the Warm Temperate/Tropical Northwestern Atlantic province. The genetic distances between these three species, recorded here, were relatively low, reflecting their recent divergence or, potentially, that they are still undergoing aa process of speciation (Rocha et al. 2007).

The geographic distribution of Narcine species is associated with the freshwater discharge of Orinoco and Amazonas, established during the Miocene (∼11 Mya) (Hoorn et al., 2017) reaching approximately 2.300 km of the Northwestern coast of South America (North Brazil Shelf province) (Rocha 2003; Floeter et al. 2008). This freshwater outflow represents one of the major biogeographic barriers in the Western Atlantic alongside the rising of Panama Isthmus (Floeter et al. 2008; Pinheiro et al. 2018; Araujo et al. 2022). The Amazon-Orinoco barrier plays a significant influence of vicariance in marine species distributed between Warm Temperate/Tropical Northwestern Atlantic ecoregions and Tropical Southwestern Atlantic ecoregion (Araujo et al. 2021; Floeter et al. 2008; Pinheiro et al. 2018; Ochoa-Zavala et al. 2024) and recently, also shows influences in speciation process in elasmobranch species (Rocha et al., 2003, 2005; Fontenelle et al., 2021; Rodrigues-Filho et al. 2023; Gales et al. 2024; Ochoa-Zavala et al. 2024).

But the influence of the Amazon-Orinoco barrier is not standardized for all organisms but in fact, is associated with different times of sedimentary depositions. For our study, we proposed a relatively lower deposition process between ∼11,8–6,8 Mya, which was followed by a sedimentary increase between ∼6,8–2,4 Mya, reaching a high deposition volume at ∼2,4 Mya (Figueiredo et al., 2009; Hoorn et al., 2017). Our results match with the beginning of sedimentation process caused by the Amazon-Orinoco plume (*Narcine* sp. x *N. bancroftii*/*N. brasiliensis*: ∼8.43 Mya; *N. bancroftii* x *N. brasiliensis* 6.29 Mya), when the deposition process was less intense which can provide a corridor to dispersal for electric rays, allowing them to surpass this barrier, and given this, disperse to Tropical Southwestern Atlantic ecoregion.

The colonization of the Atlantic Ocean from the Indian Ocean via South Africa has been well documented for a number of marine organisms, including rays (Rocha 2003; Rocha et al. 2007; Pellissier et al. 2014; Sales et al. 2019; Gales et al. 2024). Gales et al. (2024) refer to the formation of the Mid-Atlantic Barrier as a source of vicariance for the *Gymnura* species from the American and African continents. However, this barrier appears to have been highly permeable, during the past connections between the marine fishes from Brazil and the Gulf of Guinea. Sales et al. (2019) concluded that the genus *Aetobatus* reached the American coast via the Indo-Pacific/South Africa route, given that their molecular data are supported by the fact that the dorsal spots/ocelli of these rays appeared relatively recently, and are not found in *Aetobatus narutobiei* and *Aetobatus flagellum*, which are the oldest lineages, that predate the diversification of the genus in the Atlantic.

In the present study, in addition to the route mentioned above for the majority of the American narcinids (colonization of the Pacific through the East Pacific Barrier and Caribbean prior to the closure of the Isthmus of Panama), the colonization route for the genera *Benthobais*, *Discopygi*, and *Tretonarce* crossed from the Indian Ocean via South Africa to the Atlantic, and then arrived in the region of Patagonia (Magellanic realm), on the American continent, with *Benthobatis kreffti* subsequently dispersing to the southwest Atlantic and *Tetronarce tremens* to the eastern Pacific. As in other species of elasmobranch (Sales et al 2019; Hirschfeld et al. 2021; Rodrigues-Filho et al. 2023; Gales et al. 2024), biogeographic events also have influenced the dispersal and vicariance of these lineages.

### Implications for the systematics of the Narcinidae

Paraphyly of the Narcinidae and revalidation of *Syrraxis* Bonaparte (ex Jourdan) (1841) The order Torpediniformes includes 16 genera, and 72 species arranged in five families, the Torpedinidae, Hypnidae, Narcinidae, Narkidae, and Platyehinidae (Last et al. 2016). The phylogenetic relationships and systematics of these five families remain largely unclear (Moreira and Carvalho 2012; Last et al. 2016). The available studies indicate that the electric rays, except for the Paltyrhiniae, form two distinct clades within the Torpediniformes, the superfamily Torpedinoidea Jonet, 1968 and the superfamily Narcinoidea Compagno, 1973 (McEachran and Aschliman 2004; Aschliman et al. 2012; Claeson 2014; see also Compagno 1973, 1997). Two families are recognized in the Torpedinoidea – Torpedinidae Bonaparte, 1838 and Hypnidae Gill, 1862 – while the Narcinoidea is composed of the Narcinidae Gill, 1862 and Narkidae Fowler, 1934 (Carvalho, 2010; Aschlimanet al. 2012; Last et al. 2016).

In chapter 2 of their book “Rays of the World”, Last et al. (2016) do not include either the Zanobatidae or the Platyrhiniae among the principal groups of rays (Rajiformes, Torpediniformes, Rhinopristiformes, and Myliobatiformes). These authors interpret the Platyrhiniae as a sister group of the Torpediniformes, rather than a member of this order, forming a monophyletic arrangement and sister group of the Miliobatiformes and Rhinopristiformes. The results of the present study further support the exclusion of the Platyrhinidae from the Torpediniformes, given the greater phylogenetic proximity of the former with the Rajioformes, which form a single, well-supported clade (Fig. 2 and Fig.3).

The monophyly of the Narcinoidea has been discussed widely, with some studies identifying a monophyletic clade between the family Narcinidae (*Benthobatis, Diplobatis, Narcine, Discopyge*,and *Narcinops*) and the Narkidae, with the narkids being considered to have been derived from the Narcinidae (Aschlimane et al., 2012; Claeson et al., 2014; Last et al., 2016). Nayloret al. (2012) had already noted the phylogenetic proximity between the Narcinidae and the Narkidae and had recommended more detailed studies with a more comprehensive sample of the *Narcine* species. In their Chapter 2, Last et al. (2016) discuss this proximity between *Narcine* and Narkidae, and subdivide the narkids into two groups, one containing the genera *Temera* and *Narke*, and the second, *Typhlonarke*, but together with the narcinid genera *Benthobatis* and *Discopyge*. This contradicts the monophyly of the narcinid and narkid clades within the Narcinoidea. Melis et al. (2023) found a similar arrangement among *Discopyge*, *Benthobatis* and *Typhlonarke*, but diverged from the findings of Last et al. (2016) by identifying the Torpedinidae as a sister group of the genus *Narcinops*, with this Torpedinidae/*Narcinops* clade being the sister group of the *Benthobatis/Typhlonarke/Discopyge* clade. The other narkids analyzed by Melis et al. (2023) formed the sister group of this phylogenetic arrangement.

The findings of the present study indicate the existence of four narcinid clades. One clade, formed by representatives of the genera *Benthobatis* and *Discopyge*, which is the sister group of the Narkidae and Torpedinidae, while the second is the *Narcinops* clade, which is the sister clade of this first clade. The third and fourth clades contain the representatives of the genus *Narcine*, which are located in the American and Central/Western Indo-Pacific realms, respectively. While the present study, like Last et al. (2016) and Meliset al. (2023), lacked access to specimens of the other three narkid genera (*Electrolux*, *Heteronarce*, and *Typhlonarke*), the results confirm the lack of monophyly in the Narcinidae, given that three genera (*Narcinops*, *Benthobatis*, and *Discopyge*) are more closely related phylogenetically to the Narkidae and Torpedinidae.

The four clades identified here are well supported statistically, with the genera *Narcinops, Benthobatis*and *Discopyge* being recovered in clades distinct from the other narcinids. In the analysis of divergence times, these genera are phylogenetically closer to the torpedinids. Both *Narcinops*and *Benthobatis/Discopyge* present high levels of genetic distance in relation to the other narcinids (18.4–21.54 %), which are compatible with the level of differentiation between families. It is important to note here, however, that there are a number of discrepancies in comparison with two previous studies (Last et al. 2016; Melis et al. 2023), which may be related to the lack of samples of species of *Typhlonarke*, the only genus that did not group with the other narkid genera (Last et al. 2016), and the absence of *Diplobatis*.

Even so, the phylogenetic arrangement of the narkid genera that was included here (*Crossinarke* and *Narke*) is the same as that observed by Last et al. (2016). The monophyletic clade that contains the other narcinids studied here recovered a clear division of *Narcine* into two clades, from the American and Central/Western Indo-Pacific realms. These findings reflect the major genetic distances found within the genus *Narcine* (American and Indo-Pacific clades) also observed by both Last et al. (2016) and Melis et al. (2023).

Henle (1834) described the genus *Narcine* based on comparisons between European specimens of *Torpedo* and a Brazilian specimen (*Torpedo brasiliensis*, now *Narcine brasiliensis*). In particular, the frontal edge of the disc of *N. brasiliensis*, in front of the eyes, is elongated and tapered, similar to *Rhinobatos* Linck, 1790, whereas the European electric rays were rounded at the front, and in many cases, even slightly curved. In their review, Bigelow and Schroeder (1963) considered *Syrraxis* Bonaparte (ex Jourdan) (1841), *Cyclonarce* Gill (1862), *Gonionarce* Gill (1862), and *Narcinops* Whitely (1940) to be synonyms of *Narcine*. Prior to this, Gill (1862) had differentiated *Cyclonarce* and *Gonionarce* from *Narcine*, given that they both have a trilobed nasal valve, in contrast with the single lobe found in *Narcine*. *Syrraxis*, which was proposed by Jourdan and published by Bonaparte (1841) in his study of the Italian fauna, refers to a torpedine denominated *Syrraxis indica*, which was found to be no different from *Narcine indica*, described by Henle, nor any of the other species of the genus.

More recently, the Australian *Narcine* species were reallocated to *Narcinops*, a genus that was described by Whitely (1940), and was previously considered to be a synonym of *Narcine*. Last et al. (2016) considered revalidating *Narcinops* since its species are the only taxa that have a small, spade-shaped disc and an extremely elongated tail, in addition to the molecular differences observed by Naylor et al. (2012). The results of the present study further support the revalidation of the taxon *Narcinops* for the Australian electric rays, as well as the need for a review of the *Narcine* species. Given the observed separation of the American and Indo-Pacific narcinids, it is possible to confirm, based on the molecular data presented here and in other previous studies, that the genus *Narcine* is exclusive to the American littoral, as described originally by Henle (1834), based on the type species of the genus, *Narcine brasiliensis*.

In the Indo-Pacific, *Narcine indica*, currently *Narcine timlei*, is widely considered to be the type species of the three synonymous genera (*Syrraxis, Cyclonarce*and *Gonionarce*). This species is part of Central/Western Indo-Pacific clade identified in the present study. In this case, the evidence presented here reinforces the need for a thorough review and possible revalidation of the synonym *Syrraxis*, which has chronological priority as the taxon for the narcinids of the Central/Western Indo-Pacific realm.

### The N. entemedor and N. maculatalineages

One other interesting aspect of the systematics of the Narcinidae is the possible existence of distinct lineages in the rays currently identified as *N. entemedor* and *N. maculata*. Last et al. (2016) reported the occurrence of three *Narcine* species in the region of the American Pacific, *N. entemedor*, distributed between Baja California and northern Peru, *N. vermiculatus*, between the Gulf of California and Costa Rica, and *Narcine leoparda* Carvalho 2001, which is endemic to the coast of Colombia. Only *N. leoparda* was not sampled in the present study. The specimens collected in the present study from Tumbes, in northern Peru, were identified morphologically as *Narcine* cf. *entemedor*. It seems unlikely that any of these species would be identified erroneously, given that *N. entemedor* is one of the large-bodied narcinids, being close in size to the Atlantic species (*N. bancroftiii* and *N. brasiliensis*), with similar papillate spiracles, but a different pattern of ocelli. The other two species, *N. vermiculatus* and *N. leoparda*, are small-bodied narcinids, although they can be distinguished easily by the unique coloration of their spots and ocelli (Last et al. 2016).

The results of the phylogenetic analysis and species delimitation presented here nevertheless indicate that a second lineage of *N. entemedor* is found in Tumbes, which may in fact represent a new and as yet undescribed species. This species would have diverged relatively recently (∼5.55 Mya), like the *Narcine* species of the Atlantic (Fig. 4 and Table 2). This recent process of separation would also help to account for the reduced genetic distance found between the lineages of *N. entemedor* and the Atlantic species. The identification of a possible new species from Tumbes is supported by the results of the molecular analysis presented here, the reduced potential for the misidentification of the easily discriminated *N. vermiculatus* and *N. leoparda* (not sampled here), and the fact that the type locality of *N. entemedor* is Mazatlán, in western Mexico.

The taxonomic uncertainties on *N. maculata* were already well established based on previously published studies from which the sequences analyzed here were extracted. The DNA barcode analysis of Loh et al. (2023), for example, showed that the *N. maculata* lineage from the waters of Malaysia diverges clearly from all the other sequences of *N. maculata*, from India, which led them to denominate the two lineages *Narcine* cf. *maculata* sp.1and sp.2. These two lineages diverged by 4.3%, while they diverged by only 1.6% from *Narcine* cf. *oculifera* (Loh et al., 2023). While the *N. maculata* sequences analyzed here are available on the NCBI database (KF899600, KF899599, KF899598, MG644350, and MG644348), the papers that report them have not yet been formally published. The phylogenetic analyses, divergence time (∼9.37 Mya), and species delimitation presented here, together with the genetic distance between these lineages (1.9%), are all consistent with Loh et al. (2023), and reinforce the need for further, more detailed studies on *N. maculata* species complex that provides more definitive insights into the taxonomic uncertainties surrounding this species.

### Narcine bancroftii, N. brasiliensis and Narcine sp

Several studies have emphasized the importance of the variation in phenotypes in the context of both natural and sexual selection, and how the understanding of this variation can provide insights into the processes that drive the formation of populations and species (Schluter 2001; Kirkpatrick and Ravigné 2002). This variation, which is known as phenotypic plasticity, may often be the product of a prolonged process of speciation, reflecting an important adaptive strategy for the occupation of distinct environments (Herrel et al. 2001; Robinson and Parsons 2003).

Phenotypic plasticity may often be found in incipient species that have undergone recent ecological changes, providing important insights into the evolutionary mechanisms underpinning the speciation process. In general, speciation is only inferred after the event, through the comparison of closely-related species, and is thus limited to the identification of the features that differentiate the species, rather than the processes that led to their formation (McPhail 1994; Orr and Smith 1998; Beheregaray and Sunnucks 2001).

Like other narcinid and torpediniform species, *N. brasiliensis* has been reclassified a number of times, having been first described as *Torpedo brasiliensis* Olfers 1831, prior to being allocated to the genus *Narcine* Henle 1834, as *Narcine brachypleura* Miranda Ribeiro 1923, which was subsequently redescribed as *N. brasiliensis* Olfers 1831. This species was distributed across almost the entire American Atlantic coast (Miranda-Ribeiro, 1907; Bigelow and Schroeder, 1953; Figueiredo, 1977), and was divided into two subspecies – *Narcine brasiliensis corallina* Garman 1881 and *Narcine brasiliensis punctata* Garman 1881, which was posteriorly synonymized as *Narcine brasiliensis bancrofti* (Griffith, 1834)(Garman, 1913). Currently, these subspecies are synonyms of *N. bancroftii*.

Carvalho (1999) assessed specimens representing most of the distribution of *N. brasiliensis* and recognized regional morphological differences that justified its division into three species – *Narcine bancroftii*, *N. brasiliensis* and *Narcine* sp., with the latter still undescribed. This author determined that *N. bancroftii* was resident in the tropical and temperate North Atlantic Ocean, while *Narcine* sp. was restricted to the tropical Atlantic, and *N. brasiliensis* to the temperate Atlantic of South America. Other authors have questioned this arrangement, however, and have argued that the differentiation of these taxa based on morphological characters is too complex, and that the discrimination of the Atlantic species should be based primarily on their geographic distribution (Martins et al. 2009; Vianna and Vooren 2009; Rolim 2012).

These species, which were described by Carvalho (1999) based primarily on morphological and environmental characteristics, but these features, together with their history of reclassification, reflect the in numerable phenotypic variations found within the distribution of these rays along American Atlantic coast. However, Rodrigues-Filho (2020) observed low level of genetic distance between the three species – 4.02% for *Narcine sp.* x *N. bancroftii*, 2.89% for *Narcine* sp. x *N. brasiliensis*, and 1.17% for *N. bancroftii* x *N. brasiliensis* – in comparison with the mean divergence (12.54%) recorded between the other narcinids evaluated in his study. This impeded the differentiation of the three forms at the species level, based on the analysis of the DNA barcode.

In the present study, the analyses of phylogenetic inference, i.e., the Maximum Likelihood (ML), Inference Bayesiana (BI) and Neighbor-Joining (NJ), all supported the separation of the *Narcine* species from the American Atlantic, given that each species was allocated to a monophyletic group, which is consistent with the hypothesis of Carvalho (1999). However, the clades containing each species were not very well supported in all the phylogenetic analyses, which may have been due to the relatively low levels of genetic distance found between the species (*Narcine* sp. x *Narcine brasiliensis* x *Narcine bancroftii* = 0.7– 1.1%). These low genetic distances probably determined the incomplete separation of the three taxa in the species delimitation analyses.

The low *p* distances indicate that the ancestral lineage of the three forms found in the American Atlantic diverged relatively recently, which is confirmed by the divergence time analyses, that indicate that the ancestor of the three Atlantic species arrived in this region approximately 8.43 Mya, coming from the western tropical Pacific. Once this ancestral lineage had arrived in the tropical Atlantic, it likely dispersed both northward and southward, to form the current distribution and diversification of the *Narcine* species of the western Atlantic.

In particular, *N. bancroftii* is currently distributed throughout the biogeographic realms of the Temperate North and Tropical of the Atlantic (McEachran and Carvalho, 2002; Spalding et al. 2007). The former realm encompasses the temperate and subtropical waters of the North Atlantic Ocean, with the distribution of the species coinciding with the Warm Temperate Northwest Atlantic and Tropical Northwestern Atlantic provinces, where there tends to be a reduced incidence of solar radiation, typical of more temperate latitudes, with more transparent waters (O’Brien et al. 2017).

Like *N. bancroftii*, *Narcine* sp.is distributed in the Tropical Atlantic realm, albeit in the province closest to the Equator, that is, the North Brazil Shelf province. This region has much warmer water, given the hot and humid local climate (Moraes 2005), with high rainfall levels. The North Brazil Shelf province includes the world’s largest continuous tract of mangrove forest (Schaffer-Novelli et al. 1990), which is subject to the constant, intense fluvial discharge of the Amazon River and other local coastal drainage basins. This has a major influence on the physicochemical and biological characteristics of the region’s waters, reducing salinity and inserting high concentrations of suspended sediments, making the water turbid (Grodsky et al. 2014; Andrade et al. 2016). The bottom is formed predominantly by mud, derived from the input of organic matter from the rivers (Kowsmann and Costa 1979).

*Narcine brasiliensis* is also found in the tropical Atlantic realm, as well as the temperate realm of South America, predominantly in two provinces, the Tropical and the warm Temperate Southwest Atlantic provinces. These areas encompass an enormous diversity of environmental conditions, which are quite distinct from those found in the North Brazil Shelf province, where *Narcine* sp. is found. The Tropical Southwest Atlantic province is subject to far less fluvial input, with a negligible discharge of sediments, resulting in more transparent waters, influenced directly by the adjacent oceanic waters, with higher salinity and high rates of evaporation (Barreto and Summerhayes; 1975; França et al. 1976; Ekau and Knoppers 1999). By contrast, the warm temperate Southwest Atlantic province has a diversity of environments, being dominated by cooler and less saline waters resulting from the mixing of the Brazil Current with coastal waters, together with the Central South Atlantic resurgence, which brings even colder and less saline water into the area (Rossi-Wongtshcgowski et al. 2006). Both these provinces have biogenic, siliciclastic, and terrigenous carbonate sediments, associated with mangrove forests, albeit on a much smaller scale than on the North Brazil Shelf (Milliman and Summerhayes1975; Hayes 1979; Leão et al. 2003).

The low levels of genetic distance and the divergence time data indicate that *Narcine* sp., *N. brasiliensis*, and *N. bancroftii* radiated relatively recently. This evidence, together with the considerable environmental diversity found within the distribution of each species (Fig. 1; Spalding 2007), is consistent with an incipient process of diversification and speciation. Marceniuk et al. (2019) observed a similar scenario in the marine catfishes of the genus *Aspistor* resident in different areas along the Atlantic coast of South America, which were not differentiated significantly by their genetic variation, despite having clear morphological differences. These authors concluded that this morphological variation was associated with the environmental conditions faced by each species within its geographic range, given that the species were distributed from the equatorial northern coast of Brazil to the subtropical southeast, reflecting the morphological plasticity found in the species of the genus *Aspistor.* Castillo-Páez (2017) also recorded a similar scenario to that of the Atlantic *Narcine* species in the guitarfish *Zapteryx exasperata*, Jordan and Gilbert, 1880 and *Zapteryx xyster*, Jordan and Evermann, 1896 from the tropical eastern Pacific Ocean, where genetically similar individuals were distinct in their coloration.

Overall, it would seem reasonable to infer that the Atlantic *Narcine* species are subject to contrasting selective pressures, associated with the environmental differences found in the ecosystems distributed across the American Atlantic, which have driven their morphological differentiation, in particular, the variation in their coloration (Fig. 4). These morphological differences likely reflect the phenotypic plasticity associated with the distinct selective pressures that have driven diversification both within and between populations, leading to the formation of new species through adaptive radiation (Pfennig et al. 2010; Wellband and Heath 2017; Rajkov et al. 2020).

In particular, *Narcine* sp., which is found in the area dominated by the plume of the Amazon River, has the most distinct color pattern, with its dorsal surface being covered with innumerable small, dark spots of varying shape, ranging from circular to elongated, which are also found on the dorsal, pelvic, and caudal fins. In *N. bancroftii*, by contrast, the spots are much larger, dark and oval, forming cells that are rarely interconnected on the disc and the caudal peduncle, sometimes with distinct margins, either dotted or continuous. *Narcine brasiliensis*, in turn, also has large, dark oval spots that form cells, although they are sometimes interconnected on the disc and the caudal peduncle, but without the dotted margins. These varying characteristics may be associated with specific adaptations to the environmental conditions encountered by each species within its range.

The potential relationship between this phenotypic plasticity and the adaptive radiation of the electric rays likely reflects the rapid diversification of a single ancestral lineage in response to the selective pressures imposed by specific environments. This plasticity would facilitate the colonization of new environments, and the persistence of the populations, through rapid adaptation in response to selective pressures (Pfennig et al. 2010). Overall, then, the present study validates *Narcine* sp. as a lineage that is evolutionarily independent from both *N. bancroftii* and *N. brasiliensis* and emphasizes the urgent need for the formal description of the species.

## CONCLUSIONS

The present study provides valuable insights into the processes of dispersal and colonization of the oceanic environment by the ancestral lineages of the electric rays. The phylogenetic analyses, combined with the assessment of the biogeographic history of these fish permitted the estimation of the approximate colonization time of each oceanic province. These analyses confirmed the Central Indo-Pacific realm as the original center of dispersal, which supports the theory of the Tethys Sea as the origin of the region’s biodiversity, with dispersal across the Pacific through the East Pacific Barrier toward the Caribbean (Atlantic), which was colonized prior to the closure of the Isthmus of Panama, in the case of the representatives of the genus *Narcine*, and dispersal to the Atlantic through the Indian Ocean via South Africa for the representatives of the genera *Benthobatis*, *Discopyge*, and *Tetronarce*.

The data presented here indicate that the ancestor of the Torpediniformes originated at ∼153.8 Mya, which corresponds to the onset of the Cretaceous, followed by rapid radiation events in this group of rays, with the principal lineages of the families Narcinidae, Narkidae, and Torpedinidae diversifying rapidly between the end of the Cretaceous and the beginning of the Paleocene (70–60 Mya). This is consistent with the assumption that all the clades of modern batoids radiated during the lower-mid Paleogene. The present study provides evidence that the modern narcinid taxa emerged and diversified at 25–20 Mya, which is consistent with the emergence of the barriers of the Central Indo-Pacific realm, and the other diversification events mentioned above.

The results of the present study confirmed the lack of monophyly in the family Narcinidae, as well as the validity of the genus *Narcinops* as the taxon of the Australian electric rays, and the separation of the American *Narcine* species from those of the Indo-Pacific realms. In fact, the molecular data presented here, as well as in other, previous studies, confirm that the genus *Narcine* is exclusive to the American littoral, as proposed originally by Henle (1834), in his description of the type species of the genus, *Narcine brasiliensis*. In this case, the species of the Indo-Pacific would require a review and possible revalidation of the synonym *Syrraxis*, which has chronological priority, to replace *Narcine* in the Central/Western Indo-Pacific region.

The findings of the present study also revealed the existence of a second lineage of *N. entemedor* in Tumbes (Peru), which may in fact be a new and as yet undescribed species, given that the type locality of *N. entemedor* is Mazatlán, in western Mexico. A similar scenario was recorded in the case of *N. maculata*, which had distinct lineages in India and Malaysia, as found in some previous studies, reinforcing, once again, the need for further, more detailed research on *N. maculata*, in order to compile the data necessary to resolve the taxonomic uncertainties found here.

The low genetic divergence and the divergence times recorded for the *Narcine* species of the Atlantic indicate that these species diversified relatively recently, and the extensive environmental variation found within the distribution of each of the three species (*Narcine* sp., *N. brasiliensis*, and *N. bancroftii*) reflect the circumstances of an incipient process of speciation. It seems likely, in fact, that the major ecological differences found among the ecosystems distributed along the entire western Atlantic would have driven the morphological differences found among the populations (adaptive plasticity), which gave rise to the diversification and formation of the present-day species.

## Acknowledgments

We are grateful to the Phylogenomics Laboratory at the UFPA Bragança campus and the Genetics and Biotechnology Laboratory at the UFRA Capanema campus for logistic support. This study was funded by CNPq universal project number 474843/2013-0. JBLS also thanks the UFPA Institute of Biological Sciences (ICB 285/2024-ICB/UFPA) and the Shark Conservation Fund for supporting the project “Uncovering the Global Shark Meat Trade” coordinated by JBLS.

## Author contributions

Luis Fernando da Silva Rodrigues-Filho, Richard Klein Castro Silva, and Eduardo Lopes de Lima elaborated the methods employed in the present study. Luis Fernando da Silva Rodrigues-Filho, Iracilda Sampaio, Getulio Rincon, Jorge Luiz Silva Nunes, Raquel Sicca-Ramirez, and João Braullio de Luna Sales contributed to the analyses and the interpretation of the results. All the authors contributed critically to the manuscript and approved the version submitted here.

## Supplementary Information

Supplementary Table S1 - The samples and sequences used in the present study, with their respective scientific names, codes, GenBank/BOLD codes, and Biogeographic Realm or Marine Province. (A) Warm temperate Northwest Atlantic, (B) Tropical Northwestern Atlantic, (C) North Brazil Shelf, (D) Tropical Southwestern Atlantic, (E) Magellanic, (F) Tropical East Pacific, (G) Warm temperate Southeastern Pacific, (H) Arctic, (I) Central Indo-Pacific, (J) Western Indo-Pacific, (K) Temperate Australasia, (L) Temperate Northern Atlantic, (M) Temperate Southern Africa, and (N) Temperate Northern Pacific.

Supplementary Table S2 – Genetic distances between the families of the order Torpediniformes and the other rays included in the present study.

Supplementary Table S3 – Delimitation of the species of the family Narcinidae. The green bars show the species recovered in the present analysis that are consistent with the current taxonomy. The blue bars show valid species for which a single candidate species was recovered in the analysis. The yellow bars show valid species for which more than one candidate species were recovered in the analyses.

Supplementary Fig. S4 – Phylogenetic relationships among the rays, sharks, and chimeras included in the Bayesian Inference (BI) of the present study. The Posterior Probabilities (PP) are shown above the branches of each clade in the BI tree.

Supplementary Fig. S5 – Phylogenetic relationships among the rays, sharks, and chimeras included in the Maximum Likelihood (ML) analysis conducted in the present study. The SH-aLRT, aBayes, and UFBoot values are shown in the ML tree above the branches of each clade, in this order (SH-aLRT/aBayes/UFBoot).

Supplementary Fig. S6 - The estimated TMRCA and historical biogeographic reconstruction of the order Torpediniformes obtained by the BBM method, run in RASP 4.2 (Yu et al., 2020). The circle at each node represents the probability of the possible ancestral area, which is identified in the captions to the left of the tree (see Table 2 and Table S1). The estimates of divergence times are shown on the horizontal scale bar located below the tree.

## Notes

### Competing Interest Statement

The authors have declared no competing interest.

